# Evaluating single-cell cluster stability using the Jaccard similarity index

**DOI:** 10.1101/2020.05.26.116640

**Authors:** Ming Tang, Yasin Kaymaz, Brandon Logeman, Stephen Eichhorn, ZhengZheng S. Liang, Catherine Dulac, Timothy B. Sackton

## Abstract

**Motivation:** One major goal of single-cell RNA sequencing (scRNAseq) experiments is to identify novel cell types. With increasingly large scRNAseq datasets, unsupervised clustering methods can now produce detailed catalogues of transcriptionally distinct groups of cells in a sample. However, the interpretation of these clusters is challenging for both technical and biological reasons. Popular clustering algorithms are sensitive to parameter choices, and can produce different clustering solutions with even small changes in the number of principal components used, the k nearest neighbor, and the resolution parameters, among others.

**Results:** Here, we present a set of tools to evaluate cluster stability by subsampling, which can guide parameter choice and aid in biological interpretation. The R package scclusteval and the accompanying Snakemake workflow implement all steps of the pipeline: subsampling the cells, repeating the clustering with Seurat, and estimation of cluster stability using the Jaccard similarity index. The Snakemake workflow takes advantage of high-performance computing clusters and dispatches jobs in parallel to available CPUs to speed up the analysis. The scclusteval package provides functions to facilitate the analysis of the output, including a series of rich visualizations.

**Availability:** R package scclusteval: https://github.com/crazyhottommy/scclusteval Snakemake workflow: https://github.com/crazyhottommy/pyflow_seuratv3_parameter

**Supplementary information:** Supplementary data are available at *Bioinformatics* online.

## 1 Introduction

One of the most powerful applications of single-cell RNAseq is to define cell types based on the transcriptional profiles of the cells. A number of tools such as Seurat (Macosko et al. 2015), scanpy (Wolf, Angerer, and Theis 2018), and SINCERA (Guo et al. 2015) have implemented unsupervised clustering methods for single-cell RNAseq data.

Although benchmarking studies have examined the performance of different clustering algorithms (Duo et al 2018), less attention has been given to optimizing clustering algorithms for a particular dataset. Two main questions exist in this perspective. First, given a set of selected parameters, how robust is each cluster? To answer these questions, a “cluster robustness score” was proposed (Kanter, Dalerba, and Kalisky 2019). A robust cluster is defined as one that does not mix with neighboring clusters after introducing noise to the scRNAseq data. More recently, Liu et.al presented an entropy-based universal metric to measure the purity of a given single cell population (Liu et al., n.d.).

Second, what is the best way to select parameters for a specific dataset? A partial way to address this problem is clustree (Zappia and Oshlack 2018), which plots the clusters as a tree structure to visualize the relationship among clusters with different resolutions and aid in the determination of cluster numbers and appropriate parameters. However, when the cluster number is large, the tree becomes hard to interpret. Furthermore, it does not provide any quantitative assessment and requires manual inspections of the trees. An alternative, scClustViz (Innes and Bader 2018) is a Shiny app for assessing and visualizing single-cell clustering results, which uses silhouette width as a measure and the number of differential expressed genes as a rule to fine tune clustering granularity. The rationale is that the more similar clusters are, the fewer differentially expressed genes exist between them.

While these methods have begun to approach the problem, there remains an urgent need for a data-driven evaluation of the cluster stability.

## 2 Methods

To address part of the challenges of evaluating the clustering results, we deployed a re-sampling method in which we re-sample a subset of the cells from the population and repeat clustering. The cell identity is recorded for each re-sampling, and for each cluster a Jaccard index is calculated to evaluate cluster similarity before and after re-clustering. We then repeat the re-clustering for a number of times and use the mean or median of the jaccard indices as a metric to evaluate the stability of the cluster. While the theory behind this method has been extensively developed (Hennig 2007) (Lun, n.d.), practical implementations are lacking. The main existing option, the function *clusterboot* in the R package *fpc,* does not support clustering on a shared nearest neighbor (SNN) graph and is not easy to integrate with the Seurat R package. In addition, *clusterboot* is not parallelized, so it can take a very long time to re-cluster large datasets.

To address the aforementioned challenges, we implemented a snakemake (Köster and Rahmann 2012) workflow to perform the subsampling and re-clustering steps while taking advantage of multiple CPUs available in a high-performance computing cluster. In addition, we developed a new R package *scclusteval* to aid the analysis and visualization of the output from the snakemake workflow.

To facilitate the reproducibility of the workflow, we have created a docker container for the snakemake workflow which includes support for Seurat V3, respectively. The users only need to have a rds file for a Seurat object and set up the *config.ymal* file for the parameters they want to choose from and invoke snakemake through the *--use-singularity* flag. A side benefit is that the snakemake pipeline makes it easier to run a subset of the workflow. One use case is to recluster the full dataset using different parameters to evaluate clustering quality using a conventional metric such as silhouette score. After setting up the *config.ymal* for desired parameters to test, one can invoke snakemake by:

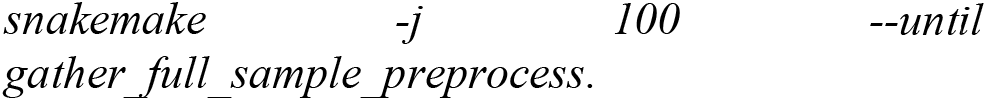

where the number after -j represents the number of CPUs to use.

The proportion of cells to be drawn at random in each subsampling (default 80%), the number of replicate samples (default 100) for the subsampling and reclustering process, and the parameters space can be configured inside the *config.ymal* file. The snakemake workflow generates two rds objects: one contains the cell identity (cluster id) information before and after the reclustering for the subsampled data and the other contains the cell identity information for the full dataset for various combinations of parameters.

The accompanying R package relies extensively on functions from the *tidyverse* packages. The fundamental object the *scclusteval* package interacts with is a tibble in a tidy format. The identity of each cell for a given set of clustering parameters is kept in a list. All the lists are kept inside the tibble as a list-column. The input of the R package is obtained directly from the snakemake pipeline output.

To explore the cell identity changes across different parameters for the full dataset, one can use the P*airWiseJaccardSetsHeatmap* function to visualize the pairwise Jaccard index across clusters (Fig1 A). Alternatively, one can use the *ClusterIdentityChordPlot* function to visualize how the cells switch from one cluster to a different cluster (Fig 1B).

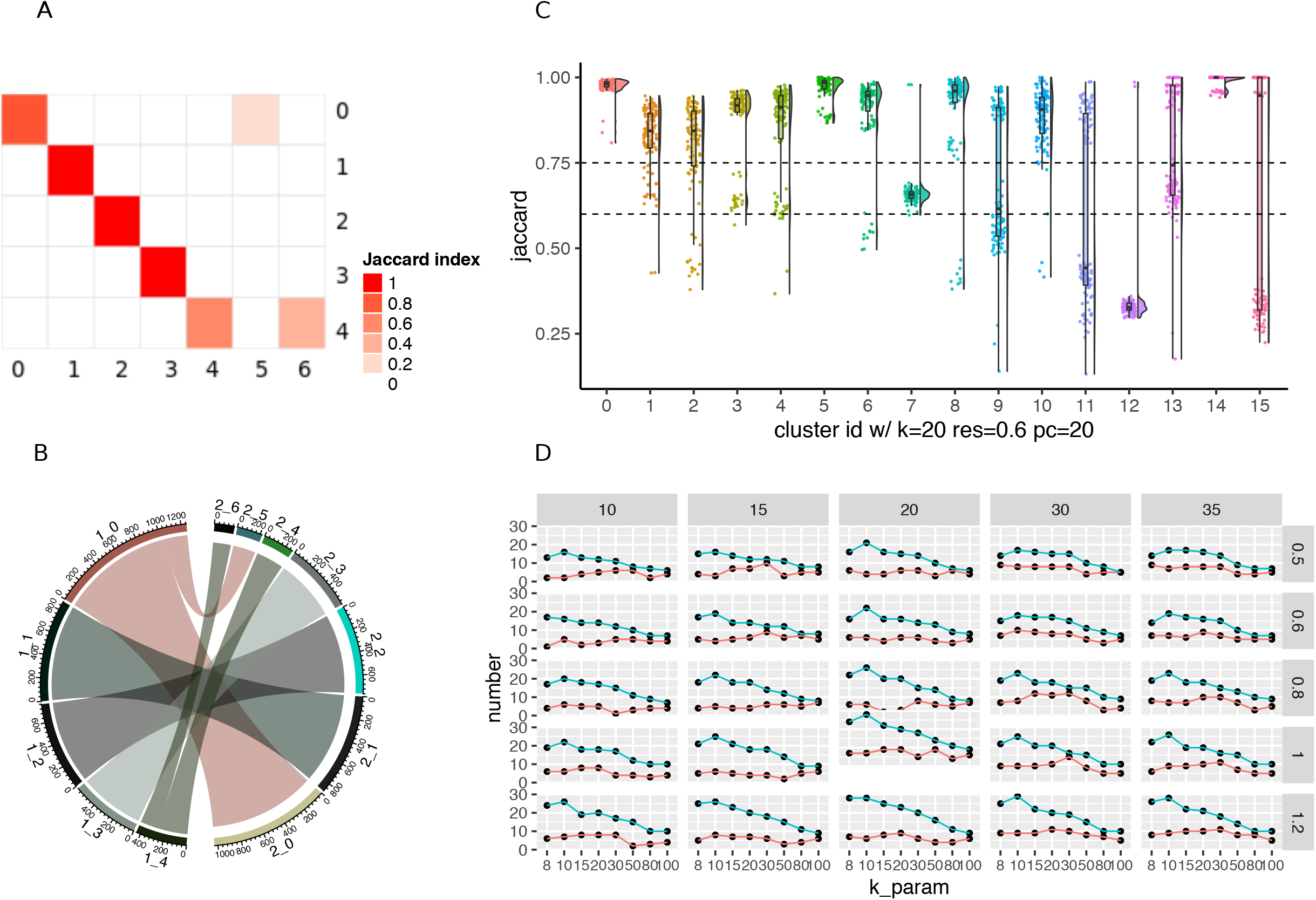
Visualizations methods from the *scclusteval* R package. (A) A pairwise Jaccard index heatmap to visualize the clusters’ relationship between two sets of different clustering parameters for the full dataset. In this example, cluster 0 in the y-axis split into cluster 0 and 5 in the x-axis; cluster 4 in the y-axis split into cluster 4 and 6 in the x-axis. (B) A cluster Chord Diagram showing cell identity switching between two different clustering parameters with additional information of the cluster size compared to (A). (C) A Jaccard Raincloud plot showing the stability of each cluster. A boxplot with a half-side violin plot showing the distribution of the Jaccard indices before and after re-clustering across 100 subsamples. (D) A line plot showing the relationship between different parameters, and the total number of clusters and number of stable clusters.

For each subset of cells, we calculate pairwise Jaccard index of each cluster before and after reclustering and assign the highest jaccard as the stability score for each cluster. The distribution of the Jaccard indices across subsamples measures the robustness of the cluster. If a cluster is robust and stable, random subsetting and reclustering will keep the cell identities within the same cluster. The heart of the visualization is the raincloud plot (Allen et al. 2019). In the raincloud plot, a boxplot together with a half-side violin plot display the distribution of the Jaccard indices. The plot can be created using the *JaccardRainCloudPlot* function. The raincloud plot gives an intuitive sense of the stability of clusters (Fig1C).

As a rule of thumb, clusters with a mean/median stability score less than 0.6 should be considered unstable. Scores between 0.6 and 0.75 indicate that the cluster is measuring a pattern in the data. Clusters with stability scores greater than 0.85 are highly stable (Zumel and Mount 2014). The *AssignStableCluster* function in our R package uses these tunable cutoffs to assign a stability level to each cluster. We observed for some datasets, the Jaccard index follows a bimodal distribution, so the mean or median may not be representative. As an alternative, we also calculate the percentage of subsampled datasets with a Jaccard index greater than a cutoff (e.g. 0.85 by default), which can be used to check stability assessments.

Because increasing resolution always generates more clusters, we also use the percentage of cells in the stable clusters to evaluate a particular clustering. We want to maximize the number of clusters but also want the majority of the cells to be in stable clusters. The *CalculatePercentCellInStable* function can be used to calculate the percentage of cells in the stable clusters.

Finally, a scatter plot to explore the relationship between the clustering parameters and the number of total clusters, total number of stable clusters, and percentage of cells in stable clusters can be made using the *ParameterSetScatterPlot* function (Fig 1D).

To demonstrate the usage of the *scclusteval* package, we analyzed two example public datasets: a mixture control dataset (Tian et al. 2019) and a 5k pbmc dataset. These analyses are available at website https://crazyhottommy.github.io/EvaluateSingleCellClustering/index.htm, powered by the workflowr (Blischak, Carbonetto, and Stephens, n.d.) R package. We also run the snakemake workflow for a mouse brain dataset and the readers can follow the tutorial to explore it. All the processed datasets can be downloaded from https://osf.io/rfbcg/.

## 3. Discussion

The graph-based clustering algorithm implemented in Seurat requires users to specify the “resolution” hyperparameter a priori. Changing it typically changes the number of clusters observed. In our analysis, increasing the resolution always produces more clusters. The set of parameters for the best resolution dependents on the biological questions under study.

Choosing the right number of PCs used for clustering is also a fundamental problem in single-cell studies. Not including a sufficient number of PCs will result in losing the true structure of the data while including too many PCs will introduce noise into the downstream analysis and may give a misleading result. A common approach is to identify the elbow point in the percentage of variance explained by successive PCs. However, this is quite subjective and not reliable. There are statistical ways to choose the number of PCs. For example, one can permutate the data matrix and calculate the eigenvalues and choose the PCs that have a higher eigenvalue than the permuted data. In general, the number of PCs scale with the complexity and heterogeneity of the data, but no general consensus on the appropriate approach to selecting PCs exists.

The k.param defines the k in the k-nearest neighbor algorithm after which a SNN graph is constructed. This parameter determines the resolution of the clustering, where a bigger k yields a more interconnected graph and larger clusters. In general, we see as the k.param goes up, the number of clusters goes down; as the resolution increases, the number of clusters goes up. The IKAP(Chen et al. 2019) tool identifies clusters by exploring the parameter space of number of PCs and k.param. The hypercluster python package uses a similar idea of exploring parameter space for clustering using Snakemake (Blumenberg and Ruggles 2020), but it is not designed specifically for scRNAseq data.

Although our Snakemake workflow was written for choosing k.param, resolution and num.pc parameters, it is flexible to extend to test other parameters. One other advantage of the Snakemake workflow is that each job is dispatched to a different CPU running in parallel and can dramatically increase the speed when testing big and heterogeneous datasets. The resulting R objects output from the snakemake are in tidy tibble format utilizing list-columns, to facilitate downstream evaluation with the *scclusteva*l R package for downstream exploration.

While the snakemake workflow is designed to interact with Seurat objects, the *scclusteval* R package can be used independently of Seurat. The input of most of the functions in *scclusteval* is a list of cell identities for a set of parameters (e.g. k_param, resolution and number of PCs) before and after reclustering. As long as the users have that in hand, *scclusteval* can be used for visualization and exploration.

We want to emphasize that finding clusters is science, but interpreting clusters is an art. There typically is not one correct clustering for any dataset, and no principled way to select a single best clustering. Note that even cell lines are not composed of pure populations of cells (Kinker et al., n.d.). There are usually cells in different cell cycle stages in typical cell cultures, and cells with different ploidies in cancer cell lines. Moreover, the concept of cell identity is evolving with the advance of the scRNAseq technology (Morris 2019; Xia and Yanai 2019). The end cluster results need to be confirmed by our understanding of the biology, and making sense of the novel clusters/cell-types/cell states is important. Our new snakemake workflow and R package provide valuable guidance in choosing parameters for clustering and facilitate the biological interpretation of the clusters derived from scRNAseq data.

## Acknowledgements

We thank

